# Estrogen promotes pro-resolving microglial behaviour and phagocytic cell clearance through the actions of annexin A1

**DOI:** 10.1101/494567

**Authors:** Rodrigo Azevedo Loiola, Edward S. Wickstead, Egle Solito, Simon McArthur

**Affiliations:** William Harvey Research Institute, Barts & the London School of Medicine & Dentistry, Queen Mary, University of London, John Vane Science Centre, Charterhouse Square, London EC1M 6BQ, United Kingdom; Laboratoire de la Barrière Hémato-Encéphalique (LBHE), Faculty Jean Perrin, EA 2465, Université d’Artois, Rue Jean Souvraz, 62307 Lens Cedex - SP18, France; School of Life Sciences, College of Liberal Arts & Sciences, University of Westminster, 115 New Cavendish Street, London W1W 6UW; institute of Dentistry, Barts & the London School of Medicine & Dentistry, Queen Mary, University of London, Blizard Institute, 4, Newark Street, London E1 2AT, United Kingdom

## Abstract

Local production of estrogen rapidly follows brain tissue injury, but the role this hormone plays in regulating the response to neural damage or in the modulation of mediators regulating inflammation is in many ways unclear. Using the murine BV2 microglia model as well as primary microglia from wild-type and annexin A1 (AnxA1) null mice, we have identified two related mechanisms whereby estradiol can modulate microglial behaviour in a receptor specific fashion. Firstly, estradiol, *via* estrogen receptor β (ERβ), enhanced the phagocytic clearance of apoptotic cells, acting through increased production and release of the protein AnxA1. Secondly, stimulation of either ERβ or the G protein coupled estrogen receptor GPER promoted the adoption of an anti-inflammatory/proresolving phenotype, an action similarly mediated through AnxA1. Together, these data suggest the hypothesis that locally produced estrogen acts through AnxA1 to exert powerful pro-resolving actions, controlling and limiting brain inflammation and ultimately protecting this highly vulnerable organ. Given the high degree of receptor selectivity in evoking these responses, we suggest that the use of selective estrogen receptor ligands may hold therapeutic promise in the treatment of neuroinflammation, avoiding unwanted generalised effects.

## Introduction

The neuroprotective potential of the steroid hormone estrogen has been the focus of numerous investigations, with epidemiological and animal model studies suggesting it may protect in conditions as diverse as stroke, Alzheimer’s disease, Parkinson’s disease and traumatic brain injury^1^. Despite this, detailed understanding of the mechanisms underlying its actions in the brain remains elusive. Many studies have focused upon the direct effects of estrogen upon damaged/dying neurones, but it is only relatively recently that the relationship between estrogen and the innate defence mechanisms of the brain, principally astrocytes and microglia, has begun to be addressed^2,3^.

Local production of estrogen by astrocytes is one of the first responses of both male and female brain tissue to injury^4^, with this production affording significant protection *in vivo*^5^. Dysregulated inflammation can be extremely deleterious for the brain^6^, hence studies have focused on suppressive actions of estradiol, revealing its ability to limit microglial iNOS activity^7^, and production of reactive oxygen species^8^, prostaglandins^9^ and inflammatory cytokines^10^.

However, it is becoming increasingly clear that in the absence of aggravating factors inflammation is naturally self-limiting, with many classical mediators activating resolving pathways^11^. This may have significant consequences for future therapeutic development, as generalized suppression of microglial activity significantly worsens neuronal loss in several animal models^12,13^, indicating the complexity of the neuroinflammatory response. We hypothesized that the actions of estrogen upon microglia are more complex than simply reducing pro-inflammatory mediator production, and that it may actively promote pro-resolving/anti-inflammatory microglial behaviour.

Microglial phagocytosis, the removal of potentially damaging threats such as invading pathogens or apoptotic cells, is central to their ability to respond to neuroinflammatory challenge. We therefore investigated the ability of estrogen to regulate microglial phagocytic clearance of apoptotic cells in unstimulated and pro-inflammatory conditions, and the consequences of estrogen treatment upon microglial phenotype, focussing on the central pro-resolving actor annexin A1.

## Materials & Methods

### Drugs

Laboratory reagents and culture media were purchased from Sigma-Aldrich (Poole, UK) unless otherwise stated. The selective estrogen receptor alpha (ERα) agonist 4,4’,4’’-(4-propyl-[1H]-pyrazole-1,3,5--triyl)trisphenol (PPT), the selective estrogen receptor beta (ERβ) agonist diarylpropionitrile (DPN), the selective G-protein coupled estrogen receptor (GPER) agonist (±)-1-[(3aR*,4S*,9bS*)-4-(6-Bromo-1,-3-benzodioxol-5-yl)-3a,4,5,9b-tetrahydro-3H-cyclop-enta[c]quinolin-8-yl]-ethanone (G1), the selective ERβ antagonist 4-[2-phenyl-5,7-bis (trifluoromethyl-)pyrazolo[1,5-a]pyrimidin-3-yl] phenol (PHTPP) and the selective GPER antagonist (3aS*,4R*,9bR*)-4-(6-Bromo-1,3-benz-odioxol-5-yl)-3a,4,5,9b-3H-cyclopenta[c]quinolone (G15) were all purchased from Tocris Bioscience, UK.

### Cell Culture

Murine microglial BV2 cells^14^ were cultured in RPMI medium supplemented with 5% fetal calf serum, 100μM non-essential amino acids, 2mM L-alanyl-glutamine and 50μg/ml gentamycin (all ThermoFisher Scientific, Poole, UK) at 37°C in 5% CO_2_. PC12 cells (ATCC) were cultured in RPMI medium supplemented with 10% normal horse serum, 5% fetal calf serum, 100μM non-essential amino acids, 2mM L-alanyl-glutamine and 50μg/ml gentamycin (all ThermoFisher Scientific, UK) at 37°C in 5% CO_2_.

### Knockdown of AnxA1 expression

BV2 cells (1□×□10^5^ cells per well) were infected with shRNA plasmids targeting murine AnxA1 from the MISSION TRC shRNA collection (Sigma-Aldrich, St. Louis, MO, USA), or with an empty plasmid control pKCON. Cells were transfected for 48h using FuGENE HD (Promega, Madison, Wisconsin, USA) according to the manufacturer’s instructions, followed by selection for stable clones using puromycin (Promega, Madison, Wisconsin, USA). Transfection was confirmed by western blot analysis.

### Phagocytosis Assay

Phagocytosis was assessed as detailed previously^15^ using 5-chloromethylfluorescein diacetate (CMFDA; Invitrogen, UK) labelled PC12 cells treated overnight with 80μM 6-hydroxydopamine hydrobromide or 40μM Na_2_S_2_O_5_ vehicle as phagocytic targets. Cells were examined using an ImageStream^×^ MKII imaging cytometer and INSPIRE software (Amnis Corporation, Seattle, WA, USA), and confirmed by parallel analysis using microscopy as described previously^15^. All experiments were performed in triplicate.

### Primary microglial cultures

Primary murine microglial cultures were prepared from 12-week-old female AnxA1 null^16^ and C57BL/6 wild-type mice, according to published protocols^17^. Cells were plated at 1.5x10^5^ cells/well in 12-well plates and cultured in DMEM supplemented with 10% FCS, 100mM nonessential amino acids, 2mM L-alanyl-glutamine, 50mg/ml gentamycin (all Life Technologies, UK) and 10ng/ml M-CSF (ThermoFisher Scientific, UK) at 37°C in 5% CO_2_ for 10 days prior to experimentation. For proliferation studies, microglia were detached from plates using accutase cell dissociation reagent (ThermoFisher Scientific, UK), prior to analysis by propidium iodide binding and flow cytometry as described below. Phagocytosis assays were performed as described above, with the additional step that cultures were immunostained for Iba1 to identify microglia as described previously^15^.

### Flow cytometry analysis

BV2 cells were fixed by incubation in 2% formaldehyde for 10 minutes prior to labelling with anti-AnxA1 50ng/ml^15^; APC-conjugated rat anti-mouse CD40 2.5μg/ml; PE-conjugated rat anti-mouse CD206 5μg/ml or appropriate isotype controls; all ThermoFisher Scientific, UK). For AnxA1 staining, cells were subsequently incubated with AF488-conjugated goat anti-mouse IgG (diluted 1:300). Inclusion of 0.025% saponin at all stages of the immunostaining process was used to define total vs. surface AnxA1 expression. In all cases, 20,000 events were acquired using a FACSCalibur flow cytometer (BD Biosciences, Cowley, UK) equipped with a 488nm argon laser; data was analysed using FlowJo 8.8.2 software (Treestar Inc, Stanford, CA, USA) with positive events being compared to appropriate secondary antibody or isotype controls.

### Quantitative RT-PCR

Total RNA was extracted using the RNeasy mini kit and genomic DNA was removed by on-column digestion using the RNase-Free DNase set, following the manufacturer’s instructions (Qiagen, Crawley, UK). cDNA was synthesized using 1μg of pooled RNA from at least three replicates using SuperScript III Reverse Transcriptase (ThermoFisher Scientific, UK). Real-time PCR was performed in triplicate, with 200ng of cDNA per well, 1μl of primers, and Power SYBR Green PCR master mix (Applied Biosystems, Warrington, UK), using the ABI Prism 7900HT sequence detection system (Applied Biosystems, UK). The following QuantiTect primers (Qiagen, UK) were used: ribosomal protein L32 (Rpl32; QT00131992), AnxA1 (QT00145915). A dissociation step was always included to confirm the absence of unspecific products. Relative expression was calculated as 2^ΔΔACT^ using Rpl32 as an endogenous control.

### ELISA

TNFα, was assayed by murine-specific sandwich ELISA using commercially available kits, according to the manufacturer’s protocols (ThermoFisher Scientific, UK). AnxA1 was assayed using an in-house ELISA as described previously^15^. Nitric oxide production was assessed using the Griess reaction for its stable proxy nitrite as previously described^15^.

### Western blot analysis

Samples boiled in 6× Laemmli buffer were subjected to standard SDS-PAGE (10%) and electrophoretically blotted onto Immobilon-P polyvinylidene difluoride membranes (Merck, UK). Total protein was quantified using Ponceau S staining (Merck, UK) and membranes were blotted for AnxA1 using an antibody raised against murine AnxA1 (1:1000; ThermoFisher Scientific, UK) in Tris-buffer saline solution containing 0.1% Tween-20 and 5% (w/v) nonfat dry milk overnight at 4°C. Membranes were washed with Tris-buffer saline solution containing 0.1% Tween-20, and incubated with secondary antibody (horseradish peroxidase–conjugated goat anti-rabbit 1:5000; ThermoFisher Scientific, UK), for 2h at room temperature. Proteins were then detected using the enhanced chemiluminescence detection kit and visualized on Hyperfilm (Amersham Biosciences, Amersham, UK). Films were digitised and analysed using ImageJ 1.51 w software (National Institutes of Health).

### Statistical Analysis

All quantified data are derived from at least three independent experiments, performed in triplicate, and are expressed as the mean ± standard error of the mean. Data were analysed by one- or two-way ANOVA as appropriate, with *post hoc* comparison using Tukey’s HSD t-test. In all cases, p ≤ 0.05 was taken as indicating statistical significance.

## Results

### Estradiol modulates microglial phagocytosis in a receptor-specific manner

Initial analysis by flow cytometry confirmed the expression of all three principal estrogen receptors (ERα, ERβ and the G-protein coupled estrogen receptor GPER) are expressed by BV2 microglia (Fig. 1A). We then investigated the ability of estrogen to modulate the phagocytosis of apoptotic PC12 cells by BV2 cells. Exposure of BV2 microglia to 17β-estradiol dose- and time-dependently enhanced the phagocytic uptake of apoptotic cells, with greatest impact after 16h treatment and at 100nM (Fig. 1B-C).

**Figure 1:**
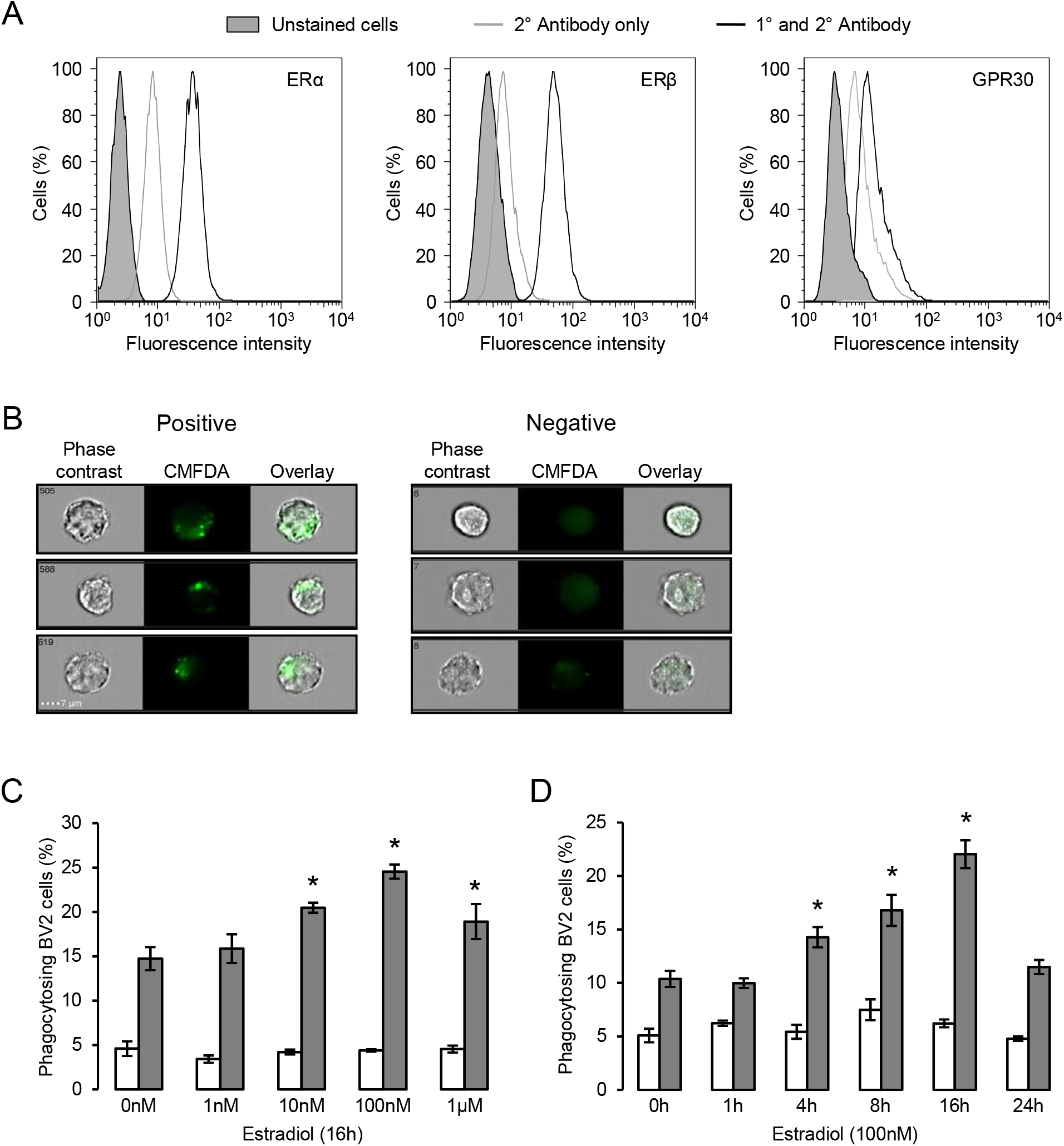
Estrogen promotes microglial phagocytosis of apoptotic cells. A) BV2 cells express all three principal estrogen receptors ERα, ERβ and GPR30, as determined by flow cytometry. Data are representative histograms of n=3 independent experiments. Exposure of BV2 cells to 17β-estradiol enhances their ability to phagocytose apoptotic (grey) but not non-apoptotic (white) PC12 cells in a B) dose and C) time-dependent manner; data are mean ± sem, n=3, *p<0.05 *vs*. untreated controls.

To determine which of the three estrogen receptors were responsible for the pro-phagocytic effects of estradiol, we examined the effect of treatment with specific pharmacological agonists. Treatment of BV2 cells for 16h with the ERα selective agonist PPT was without effect (Fig. 2A) but treatment with the ERβ agonist DPN significantly and dose-dependently enhanced phagocytosis (Fig. 2B). In contrast, 16h exposure to the GPER agonist G1 dose-dependently attenuated the ability of microglia to phagocytose apoptotic cells (Fig. 2C). Furthering this analysis, we studied the effects of the ERβ antagonist PHTPP or the GPER antagonist G15 (each administered at their respective IC_50_ values) upon the response of BV2 cells to estradiol. Pre-treatment with either antagonist alone had no effect on phagocytosis of apoptotic cells, but administration of PHTPP almost completely inhibited the pro-phagocytic effect of estradiol (Fig. 2D). In contrast, pre-treatment with G15 significantly potentiated apoptotic cell clearance by BV2 cells treated with estradiol (Fig. 2E).

**Figure 2:**
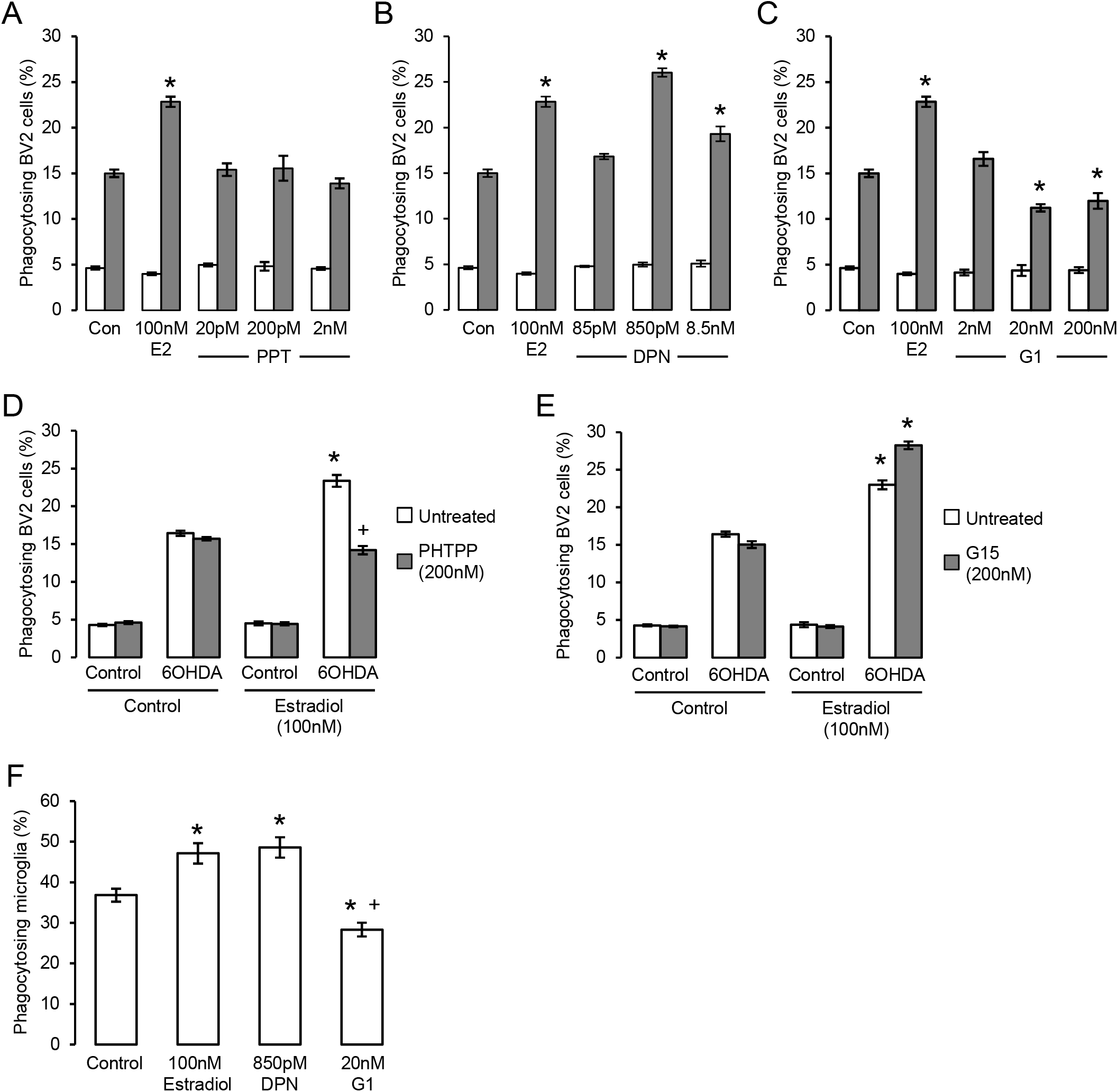
Estrogen has receptor specific effects upon microglial phagocytosis. AC) Treatment of BV2 cells for 16h with selective agonists for A) ERα (PPT), B) ERβ (DPN) or C) GPER (G1) revealed receptor dependent effects upon microglial phagocytosis, with activation of ERβ promoting and GPER inhibiting phagocytic activity; ERα activation had no effect on phagocytosis; data are means ± sem, n=3, *p<0.05 *vs*. untreated controls. D) Pre-treatment of microglia with the selective ERβ antagonist PHTPP (200nM, 15 minute pre-treatment) prevented the stimulatory effect of estradiol (100nM, 16h) upon microglial phagocytosis of apoptotic cells; data are means ± sem, n=3, *p<0.05 *vs*. untreated controls, +p<0.05 *vs*. 17β-estradiol treatment. E) Pretreatment of microglia with the selective GPER antagonist G1 (200nM, 15 minute pretreatment) enhanced the stimulatory effect of estradiol (100nM, 16h) upon microglial phagocytosis of apoptotic cells; data are means ± sem, n=3, *p<0.05 *vs*. untreated controls, +p<0.05 *vs*. 17β-estradiol treatment. F) Treatment of primary murine microglia for 16h with 17β-estradiol or DPN enhanced phagocytosis of apoptotic cells, whilst treatment with G1 inhibited phagocytosis; data are means ± sem, n=3, *p<0.05 *vs*. untreated controls, +p<0.05 *vs*. 17β-estradiol treatment.

To validate the pro-phagocytic effects of estrogen, we examined whether primary microglia derived from adult mouse brain would respond similarly to estradiol or its receptor-specific mimetics. Primary microglia were significantly more efficient than BV2 cells at phagocytosing apoptotic PC12 cells, but they nonetheless responded in a similar way to estrogen receptor ligand treatment, with estradiol or DPN treatment both potentiating phagocytosis and G1 treatment impairing it (Fig. 2F). Together, these data strongly indicate that estradiol exerts dual, and opposing, effects upon microglial phagocytosis acting via both ERβ and GPER.

### Estradiol promotes microglial phagocytosis through the mobilization of the pro-phagocytic factor, annexin A1

We have previously shown the microglia-secreted protein annexin A1 (AnxA1) to serve as an important “eat me” signal for phagocytosis following its release from microglia and subsequent binding to exposed phosphatidylserine on the apoptotic cell surface^15^. As we and others have shown this protein to mediate some of the actions of estradiol in other contexts^18,19^, we hypothesized that mobilization of AnxA1 may underlie the ERβ-dependent pro-phagocytic effects of estradiol. We therefore compared the effects of the hormone upon phagocytic behaviour of primary microglia derived from adult wild-type and AnxA1 null mice. Pre-treatment of primary microglia from wild-type animals with estradiol significantly increased apoptotic cell phagocytosis as expected, but this was not the case in microglia from AnxA1 null mice, where estradiol not only failed to augment microglial phagocytosis, but actually inhibited it (Fig. 3A).

**Figure 3:**
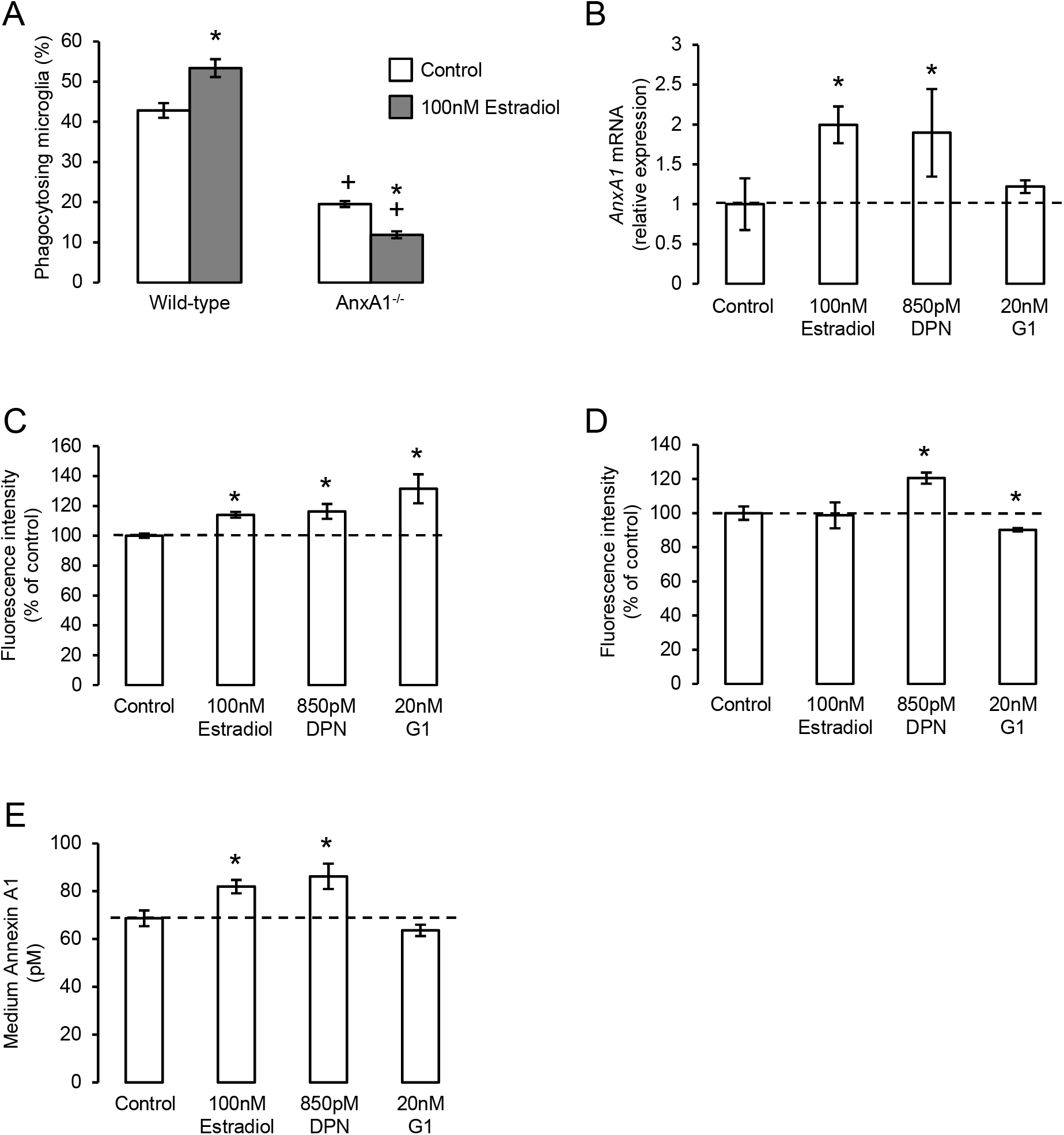
The pro-phagocytic effects of estrogen via ERβ require mobilization of annexin A1. A) Treatment of primary microglia from wild-type mice with 17β-estradiol (100nM, 16h) promotes phagocytosis of apoptotic PC12 cells, whereas similar treatment of AnxA1^-/-^ primary microglia causes a significant further inhibition in their phagocytic ability; data are means ± sem, n=3, *p<0.05 *vs*. untreated controls, +p<0.05 *vs*. similarly treated wild-type group. B) Treatment of BV2 cells for 16h with 17β-estradiol or the ERβ agonist DPN, but not the GPER agonist G1, enhances production of *AnxA1* mRNA; data are means ± sem, n=3, *p<0.05 *vs*. untreated controls. C) Treatment of BV2 cells for 16h with 17β-estradiol, DPN or G1 enhances total cellular AnxA1 content; data are means ± sem, n=3, *p<0.05 *vs*. untreated controls. D) Treatment of BV2 cells for 16h with DPN enhances, whilst similar treatment with G1 reduces, surface expression of AnxA1; data are means ± sem, n=3, *p<0.05 *vs*. untreated controls. E) Treatment of BV2 cells for 16h with 17β-estradiol or DPN but not G1 enhances release of AnxA1 into the culture medium; data are means ± sem, n=3, *p<0.05 *vs*. untreated controls.

Given the importance of AnxA1 secretion from microglia for efficient phagocytosis, we investigated how treatment with estrogen or its receptor-specific mimetics would affect the cellular localisation of the protein. Exposure of BV2 cells to either estradiol or the ERβ agonist DPN increased AnxA1 mRNA and total protein content (Fig. 3B, C), and in the case of DPN also increased cell surface AnxA1 expression (Fig. 3D). Moreover, both ligands significantly induced protein release into the culture medium (Fig. 3E). In contrast, treatment with the GPER agonist G1 had no significant effect upon AnxA1 mRNA (Fig. 3B) or secreted protein (Fig. 3E), but significantly enhanced total AnxA1 content (Fig. 3C) whilst decreasing cell surface protein (Fig. 3D), strongly suggesting increased intracellular accumulation of the protein.

### Estradiol regulates inflammatory microglial activation, promoting resolution

Outside of development and certain defined areas of the brain, non-phlogistic neuronal apoptosis is rare, with cell death being more commonly associated with neuroinflammation and disease^20^. We therefore investigated the effects of estradiol upon microglia under inflammatory conditions, modelled by exposure to bacterial endotoxin^15,21^. As previously described^15^, pre-exposure of BV2 cells to 50ng/ml LPS for 18h significantly increased the inappropriate phagocytosis of non-apoptotic cells, but this was markedly attenuated by subsequent (2h after LPS stimulation) addition of estradiol or DPN, but not G1, indicative of an ERβ-mediated action (Fig. 4A). Similarly, LPS-induced production of intracellular reactive oxygen species was reduced by treatment with estradiol and DPN, but not G1 (Fig. 4B). In contrast, estradiol, DPN and G1 were all able to attenuate LPS-induced production of the inflammatory mediators TNFα and nitric oxide (Fig. 4C, D). Characterization of BV2 cell surface markers of inflammatory phenotype revealed that treatment with estradiol, DPN or G1 could attenuate LPS-induced expression of the pro-inflammatory marker CD40 (Fig. 4E, typical profiles Supplementary Fig. 1A) and prevent LPS-induced suppression of the anti-inflammatory marker CD206 (Fig. 4F, typical profiles Supplementary Fig. 1B). Together these data suggest that estrogen receptor activation could limit the pro-inflammatory stimulation of microglia, with significant roles for both ERβ and G1.

**Figure 4:**
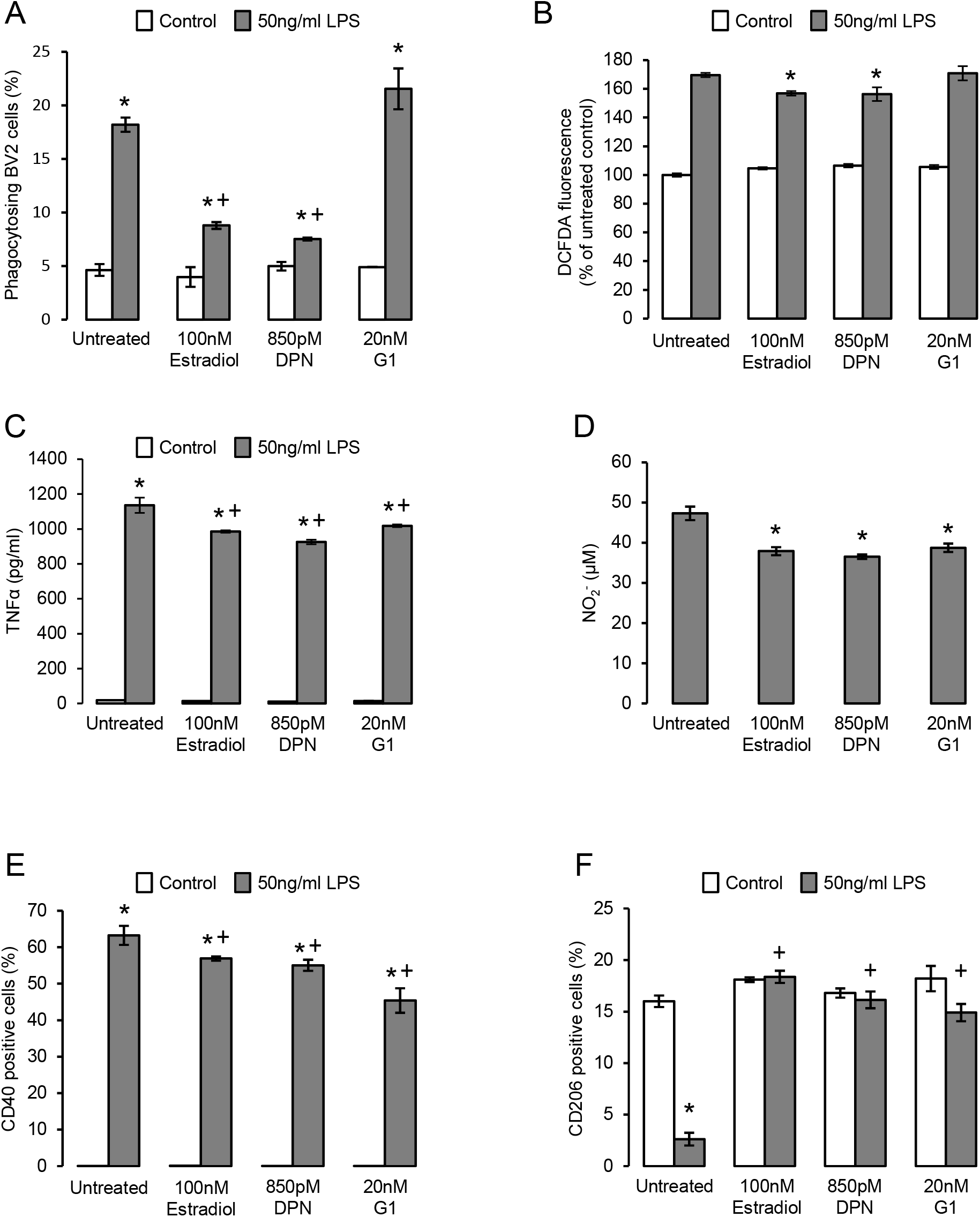
Estrogen promotes an anti-inflammatory microglial phenotype. A) Pretreatment of BV2 cells for 2h with 50ng/ml bacterial lipopolysaccharide (LPS) induces the phagocytosis of non-apoptotic PC12 cells, an effect reversed by subsequent treatment (16h) with either 17β-estradiol or DPN, but not G1; data are means ± sem, n=3, *p<0.05 *vs*. respective control cells, +p<0.05 *vs*. LPS treatment alone. B) Treatment of BV2 cells with 17β-estradiol or DPN but not G1 (16h) suppresses LPS-induced reactive oxygen species production (2h pre-treatment); data are means ± sem, n=3, *p<0.05 *vs*. LPS-treated controls. C) Treatment of BV2 cells (16h) with 17β-estradiol or DPN but not G1 (16h) limits LPS-induced production of the pro-inflammatory cytokine TNFα (2h pre-treatment); data are means ± sem, n=3, *p<0.05 *vs*. respective control cells, +p<0.05 *vs*. LPS treatment alone. D) Treatment of BV2 cells with 17β-estradiol, DPN or G1 (16h) suppresses LPS-induced production of nitrite (2h pre-treatment; baseline nitrite production was below detection limits); data are means ± sem, n=3, *p<0.05 *vs*. LPS-treated control cells. E) Treatment of BV2 cells with 17β-estradiol, DPN or G1 (16h) suppresses LPS-induced surface expression of the pro-inflammatory marker CD40 (2h pre-treatment); data are means ± sem, n=3, *p<0.05 *vs*. untreated control cells, +p<0.05 *vs*. LPS-treated control cells. F) Treatment of BV2 cells with 17β-estradiol, DPN or G1 (16h) reverses the LPS-induced loss of surface expression of the antiinflammatory marker CD206 (2h pre-treatment); data are means ± sem, n=3, *p<0.05 *vs*. respective control cells, +p<0.05 *vs*. LPS treatment alone.

Given the major pro-resolving and anti-inflammatory functions of AnxA1 identified from studies of other immune cells^22^, and our previous detection of a key mediating role for this protein in estrogen-induced phagocytosis, we hypothesised that it may similarly mediate the anti-inflammatory actions of estrogen and its receptor mimetics. We therefore stably transfected BV2 cells with a lentiviral vector bearing an shRNA sequence targeting AnxA1 to reduce expression of the protein (Supplementary Fig. 2) and investigated whether the anti-inflammatory effects of estrogen or its mimetics were maintained. Knock-down of AnxA1 in this way significantly impaired the ability of estradiol, DPN or G1 to reverse LPS-induced TNFα release (Fig. 5A), but did not affect the inhibitory effects of estradiol, DPN or G1 on LPS-induced nitric oxide release (Fig. 5B). Suppression of AnxA1 expression did however, reverse the effects of estradiol, DPN or G1 on both CD40 (Fig. 5C) and CD206 (Fig. 5D) expression, confirming an important role for AnxA1 in mediating many, although not all, of the anti-inflammatory actions of estrogen.

**Figure 5:**
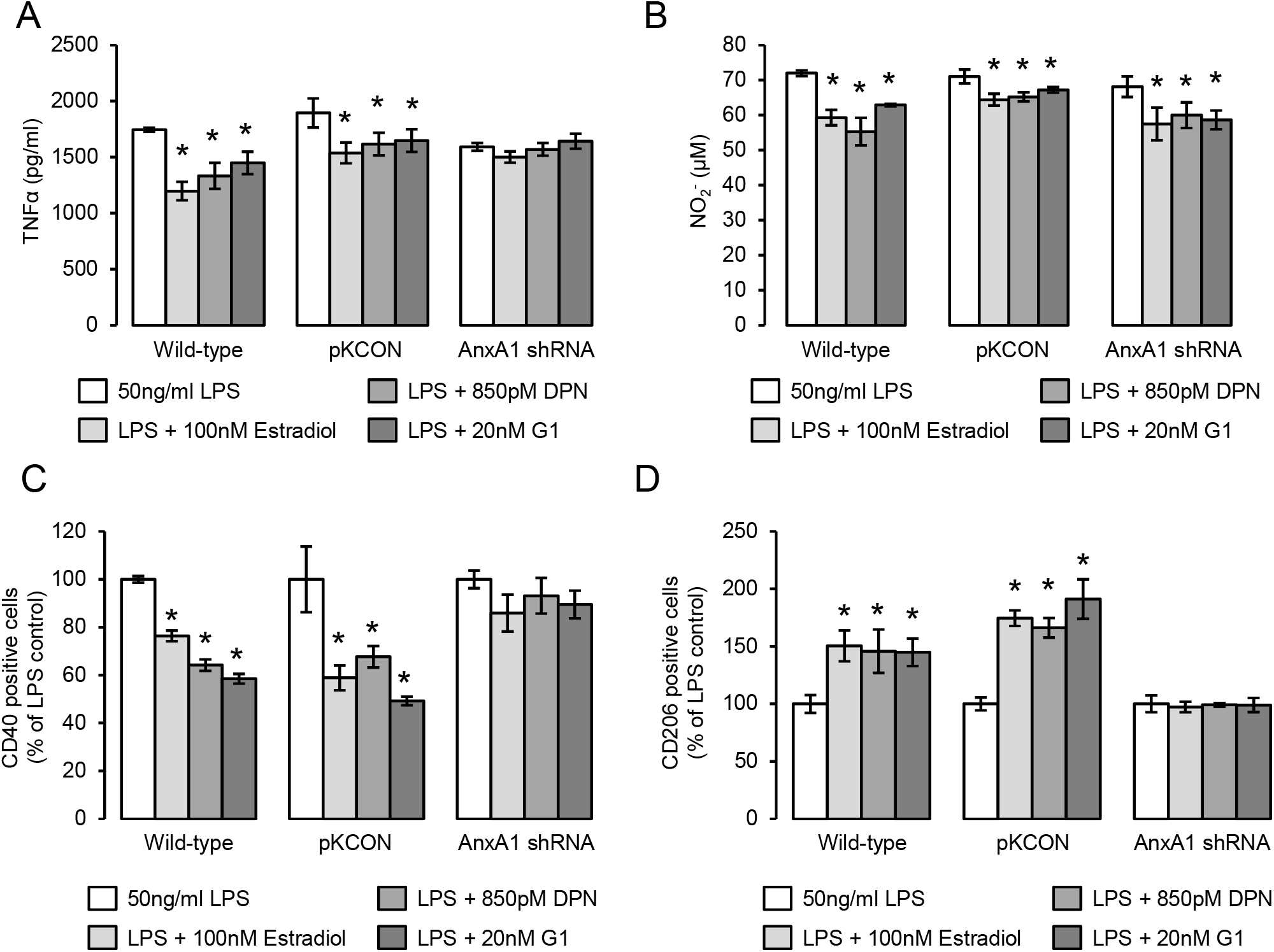
The anti-inflammatory effects of estrogen are largely dependent upon AnxA1 expression. A) Stable transfection of BV2 cells with an shRNA sequence targeting AnxA1, but not with an empty plasmid control, inhibits the ability of estradiol, DPN or G1 (16h) to suppress LPS-induced TNFα production (2h pre-treatment); data are means ± sem, n=3, *p<0.05 *vs*. LPS-treated control cells. B) Neither stable transfection of BV2 cells with an empty plasmid control nor with an shRNA sequence targeting AnxA1 affects the ability of estradiol, DPN or G1 (16h) to suppress LPS-induced nitrite production (2h pre-treatment); data are means ± sem, n=3, *p<0.05 *vs*. LPS-treated control cells. C) Stable transfection of BV2 cells with an shRNA sequence targeting AnxA1, but not with an empty plasmid control, inhibits the ability of estradiol, DPN or G1 (16h) to suppress LPS-induced surface expression of the pro-inflammatory marker CD40 (2h pre-treatment); data are means ± sem, n=3, *p<0.05 *vs*. LPS-treated control cells. D) Stable transfection of BV2 cells with an shRNA sequence targeting AnxA1, but not with an empty plasmid control, inhibits the ability of estradiol, DPN or G1 (16h) to reverse LPS-suppressed expression of the anti-inflammatory marker CD206 (2h pre-treatment); data are means ± sem, n=3, *p<0.05 *vs*. LPS-treated control cells.

## Discussion

Estrogen is increasingly recognised as a powerful modulator of immune cell activity, able to regulate cells of both the innate^23^ and adaptive^2^ arms of the immune system, actions which contribute to the sex differences commonly seen in inflammatory disorders. Inflammatory diseases of the CNS, including such major conditions as Alzheimer’s disease, Parkinson’s disease and multiple sclerosis, have similarly been shown to be sexually dimorphic in incidence and/or symptom severity^24,25^. Whilst multiple factors undoubtedly contribute to these sex differences, direct neuroprotective effects of estrogen are strongly supported^26,27^. For many years, studies have focussed on the directly neuroprotective actions of estrogen upon neurones^28^, but more recent attention has been given to the modulatory effects of the hormone upon microglial behaviour, given the importance of these cells in many neurological disorders^21,29–31^. In this study, we have identified the ability of estrogen to modulate a key aspect of microglial function in disease, the removal of apoptotic cells. We describe a clear, receptor-specific, modulatory action of the principal estrogen 17β-estradiol upon microglial clearance of apoptotic cells, and highlight the ability of the steroid to suppress inflammatory microglial activation. Moreover, we reveal a central role for the powerful pro-resolving protein AnxA1 in mediating both the pro-efferocytosis and anti-inflammatory effects of estrogen upon microglia.

Clearance of apoptotic cells by phagocytosis is critical for the maintenance of healthy tissue, as apoptotic cells that are not removed will progress to secondary necrosis and become significant inflammatory foci themselves^32^. Microglia are the principal phagocytes of the CNS and play a central role in this process, a particularly important action given the susceptibility of neurones to inflammatory damage^33^. Our data reveal estrogen to strongly potentiate microglial phagocytosis of apoptotic cells, acting through induction of AnxA1 expression and secretion. We have previously shown AnxA1 to induce a non-phlogistic phenotype in microglia post-phagocytosis^15^, strongly suggesting that its modulation by estrogen is a significant component of the hormone’s neuroprotective effects. Previous work has suggested that estrogen can promote phagocytosis by peripheral macrophages^34,35^, acting primarily through ERα. In contrast, whilst we identified ERα expression in microglia, activation of this receptor with the specific ERα agonist PPT had little impact on phagocytosis, with much clearer roles for the alternative estrogen receptors ERβ and GPER.

Whilst the overall effect of 17β-estradiol was to stimulate microglial phagocytosis, pharmacological analysis of the specific receptors involved in this process revealed an intriguing distinction between ERβ and GPER, with ERβ acting to promote phagocytosis and GPER inhibiting it. The mechanism underlying this effect appears to revolve around differential effects upon AnxA1 localisation; ERβ activation increased AnxA1 synthesis and release, whilst GPER stimulation led to intracellular AnxA1 accumulation and a reduction in secreted protein. The pro-phagocytic actions of AnxA1 are dependent upon its secretion from microglia and consequent binding to phosphatidylserine on apoptotic target cells^15^, hence the negative effects of GPER stimulation upon AnxA1 release presumably underlies the effects of this receptor upon phagocytosis. Interestingly, this dichotomy of effect was not replicated in the effects of estrogen receptor activation upon microglial phenotype, with both ERβ and GPER selective ligands being equipotent in ameliorating LPS-induced pro-inflammatory signs. This finding accords well with published studies that show exogenous administration of the GPER agonist G1 to prevent inflammatory microglial activation *in* vivo^36–38^, and that identify a mediatory role for GPER in the anti-inflammatory effects of estrogen in cerebral ischaemia^39^. These anti-inflammatory effects of estrogen or its receptor-specific mimetics again appeared to be mediated in large part through AnxA1, with microglia stably bearing AnxA1-targeting shRNA sequences showing a clear impairment in many of the anti-inflammatory actions of estrogen and its mimetics.

Our findings reinforce the importance of AnxA1 in the nervous system response to injury^40^, with microglia lacking AnxA1 being severely impaired in their phagocytic ability. Moreover, such microglia show a markedly altered response to estradiol with exposure to the hormone actually inhibiting phagocytosis, and inducing only limited antiinflammatory actions. These data thus reinforce the complexity of the systems that have evolved to regulate microglial function. Moreover, this work highlights the potent regulatory action of estrogen upon AnxA1 expression and activity^18,41^, and extends this role to the cells of the CNS. Thus, complementing its long-studied role as a mediator of glucocorticoid immunomodulatory action^22^, our results further emphasise the importance of AnxA1 in the anti-inflammatory effects of female sex hormones.

Local up-regulation of aromatase expression and consequent estrogen production is a key part of the response to CNS injury^4,5,42^, acting to limit damage and preserve tissue integrity. Where and how estrogen exerts its protective effects is less clear however, but blockade of estrogen production by genetic deletion of aromatase leads to significantly enhanced post-injury gliosis^43^. The role of microglia in such injury-induced gliosis is complex, as these cells show dramatic and dynamic changes in phenotype^44^, and have the potential to be both harmful and protective to surrounding neurones^45^. Our findings suggest that a major action of estrogen upon microglia is to promote their more protective functions, enhancing the efferocytosis of damaged/dying cells and limiting expression of pro-inflammatory features. These actions are supported by numerous studies showing exogenous estrogen to be protective in models of neuroinflammatory/neurotoxic damage^27,46^, and particularly by previous work identifying estrogen as an anti-inflammatory modifier of microglial phenotype^47–53^. Together with these reports, our study places estrogen as a significant pro-resolving mediator within the brain, acting to stimulate the removal of cellular debris post-injury and to promote an anti-inflammatory environment, highlighting its role as a major part of the brain’s endogenous defence mechanisms.

## Supporting information

## Acknowledgements

This work was supported by Alzheimer’s Research UK Pilot Grant ARUK-PPG-2016B to SM, and by FISM Fondazione Italiana Sclerosi Multipla (cod 2014/R/21) and Alzheimer’s Research UK Pilot Grant ARUK-PPG2013B to ES; ESW is supported by a PhD Scholarship from the University of Westminster.

## Author Contribution Statement

RA, ESW and SM performed experiments and analysis, SM conceived and designed the study, ES provided valuable insight and advice throughout the project. All authors contributed to the writing of the final manuscript.

## Data Sharing

No datasets were generated or analysed during the current study.

