## Supplementary material for "Estrogen promotes pro-resolving microglial behaviour and phagocytic cell clearance through the actions of annexin A1"

**Supplementary Information**

**Supplementary Figure 1:** Typical surface expression profiles for A) CD40 and B) CD206 on BV2 cells treated for 16h with estradiol (100nM) , DPN (850pM) or G1 (20nM) ± 2h prior treatment with LPS (50ng/ml).

**Supplementary Figure 2:** Western blot analysis of AnxA1 expression in three independent samples of wild-type BV2 cells or BV2 cells stably transfected with a pKCON empty plasmid or the same plasmid bearing an shRNA sequence for AnxA1. Comparison is made with total loaded protein, as visualised by Ponceau S staining.

Supplemental Figure 1

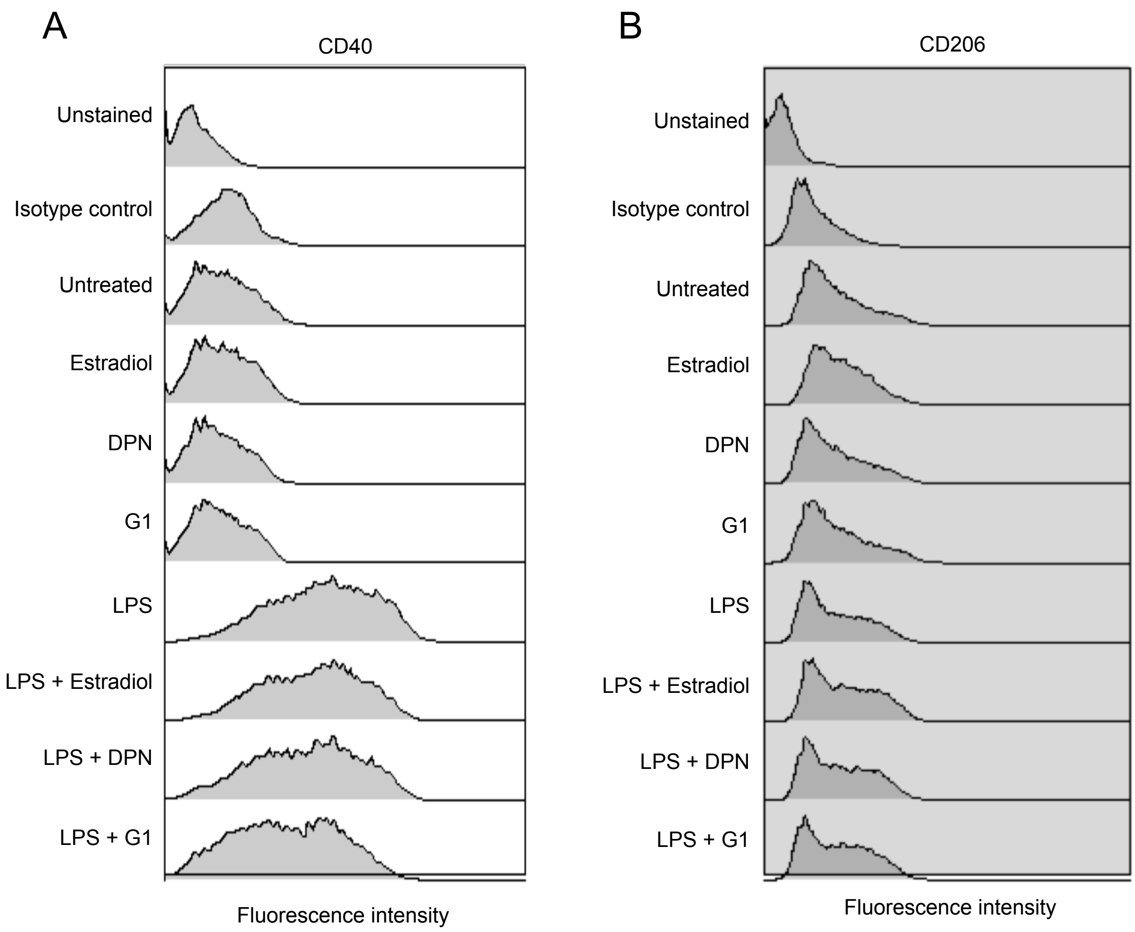

Supplemental Figure 2

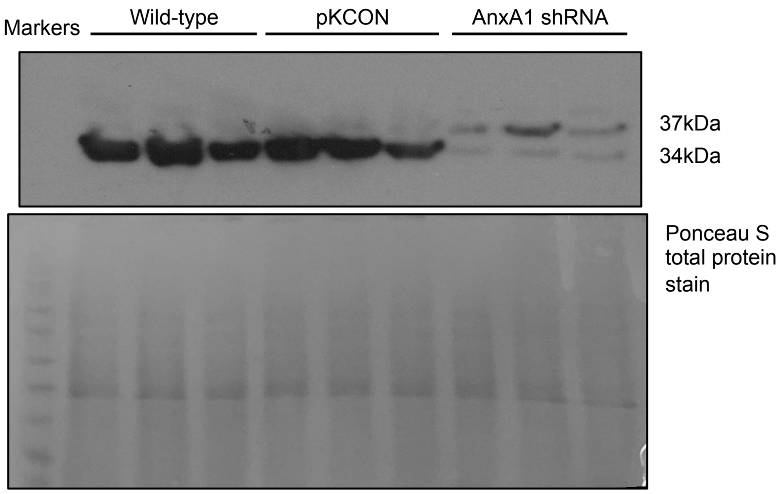
